# Combinatorial *in vivo* genome editing identifies widespread epistasis during lung tumorigenesis

**DOI:** 10.1101/2024.03.07.583981

**Authors:** Jess D. Hebert, Yuning J. Tang, Laura Andrejka, Steven S. Lopez, Dmitri A. Petrov, Gábor Boross, Monte M. Winslow

**Author notes:** Corresponding authors: Monte M. Winslow, Stanford University School of Medicine | 279 Campus Drive, Beckman Center B256, Stanford, CA 94305 | | |, Gábor Boross, Institute of Evolution | Konkoly-Thege Miklós út 29-33, Budapest, Hungary, 1121, Dmitri Petrov, Stanford University Department of Biology | 327 Campus Drive, Bass Biology 232, Stanford, CA 94305 |.

## Abstract

Lung adenocarcinoma, the most common subtype of lung cancer, is genomically complex, with tumors containing tens to hundreds of non-synonymous mutations. However, little is understood about how genes interact with each other to enable tumorigenesis *in vivo*, largely due to a lack of methods for investigating genetic interactions in a high-throughput and multiplexed manner. Here, we employed a novel platform to generate tumors with all pairwise inactivation of ten tumor suppressor genes within an autochthonous mouse model of oncogenic KRAS-driven lung cancer. By quantifying the fitness of tumors with every single and double mutant genotype, we show that most tumor suppressor genetic interactions exhibited negative epistasis, with diminishing returns on tumor fitness. In contrast, *Apc* inactivation showed positive epistasis with the inactivation of several other genes, including dramatically synergistic effects on tumor fitness in combination with *Lkb1* or *Nf1* inactivation. This approach has the potential to expand the scope of genetic interactions that may be functionally characterized *in vivo*, which could lead to a better understanding of how complex tumor genotypes impact each step of carcinogenesis.

## INTRODUCTION

Genetic interactions, in which the phenotype driven by a genetic alteration depends on the underlying genotype, indicate functional relationships between genes. Genetic interactions underlie all aspects of development, homeostasis, and disease, with most human traits being under complex genetic control^1–4^. Nevertheless, most mechanistic studies alter only one or a few genes at a time, limiting their ability to explore how gene products and pathways interact within complex biological settings. Systematic genetic interaction maps in yeast and in mammalian cell lines have revealed a multitude of functional insights into many aspects of biology^5–10^. For instance, maps of these interactions have allowed rapid identification of new functional complexes, predicted roles for uncharacterized genes, revealed unexpected network rewiring in response to environmental changes, and demonstrated functional repurposing of complexes and interactions during evolution^5,6,8–13^.

Cancer genome sequencing has cataloged somatic alterations at the genome-wide level and identified many putative driver genes^14,15^. However, identifying recurrent genomic alterations does not necessarily reveal their functional importance to cancer growth, and the impact of gene inactivation remains difficult to glean from cancer genome sequencing data alone^16–18^. Tumor suppressor gene alterations in particular are diverse and have non-random patterns of co-occurrence, suggesting underlying genetic interactions^14,19–21^. Moreover, the frequency of the rates of co-occurrence cannot generally determine how biologically important inactivation of a gene is within the context of the other alterations within a tumor^19^.

The impact of combinatorial tumor suppressor gene inactivation on neoplastic growth has been investigated using high-throughput studies on cell lines and in much lower throughput using mouse models^22–26^. Cancer cell lines have limited ability to provide insight into how tumor suppressor genes interact to constrain the expansion of tumors *in vivo* due to their near-optimal growth in culture, their widespread genetic and epigenetic changes, and the lack of an autochthonous microenvironment^18^. Conversely, genetically engineered mouse models enable the introduction of defined alterations in oncogenes and tumor suppressor genes in normal somatic cells. The resulting tumors grow entirely within their natural *in vivo* setting^27–30^. The analysis of combinatorial genetic alterations on tumorigenesis *in vivo* using conventional Cre/Lox-based systems has uncovered important genetic interactions between tumor suppressor genes^31^. However, conventional *in vivo* cancer models are not sufficiently scalable to generate a broad understanding of genetic interactions between tumor suppressor genes that drive carcinogenesis^18,27,28,32^.

CRISPR/Cas9-mediated somatic genome editing in somatic cells has increased the scale of *in vivo* functional analyses, and integration with tumor barcoding has enabled precise and multiplexed quantification of tumor initiation and growth^23,33–35^. We have generated moderate-scale maps of gene function within autochthonous cancer models by inactivating many genes in parallel in mouse models of lung cancer using pools of barcoded sgRNA-containing lentiviral vectors. Tumor barcoding coupled with high-throughput barcode sequencing (Tuba-seq) can quantify the size of each barcoded clonal tumor and the number of tumors of each genotype, enabling precise measurement of the impact of each gene on tumorigenesis^34^. Epistatic interactions between tumor suppressor genes during cancer growth *in vivo* can be assessed by combining Cre/Lox and CRISPR/Cas9 genome editing^22^. However, the requirement for a floxed allele, the considerable time and effort to generate the required compound mouse strains, and cross-cohort comparisons make it unlikely that an accurate and comprehensive analysis of pairwise interactions will be achieved in this manner.

Here we establish and validate an efficient platform, as well as associated analytic and statistical methods, to integrate Tuba-seq with combinatorial CRISPR/Cas9-mediated gene inactivation within autochthonous mouse cancer models. To uncover the broad epistatic interactions that drive lung tumor initiation and growth *in vivo*, we initiated lung tumors with a pool of barcoded dual-sgRNA vectors and quantified the effect of pairwise inactivation of every combination of ten tumor suppressor genes. Quantitative and multiplexed analysis of the combinatorial effects of cancer drivers within autochthonous models can uncover the epistatic interactions of these genes and pathways, enable a better understanding of cancer evolution, illuminate novel aspects of tumorigenesis, and potentially highlight new therapeutic opportunities.

## RESULTS

### Generation of barcoded lentiviral libraries for dual sgRNA screening with clonal resolution

Cellular barcoding has facilitated the quantification of clonal heterogeneity, and the integration of clonal barcoding into CRISPR screens increased resolution both *in vitro* and *in vivo*^34,36^. Viral vectors with pairs of sgRNAs have been used for combinatorial genome editing of cells in culture, but these have lacked the ability to provide simultaneous clonal information^37,38^. Conversely, while barcoded dual-sgRNA vectors have led to key insights into the tumorigenic potential of complex tumor genotypes *in vivo*, they have required laborious individual cloning^39–42^ and/or have suffered from lentiviral template switching that decouples the barcode from the sgRNA^43^. To generate barcoded lentiviral-sgRNA vector pools for tumor barcoding and high-throughput barcode sequencing (Tuba-seq), we recently integrated a diverse barcode within the 21 nucleotides of the U6 promoter directly adjacent to a single sgRNA (U6 barcode Labeling with per-Tumor Resolution Analysis; Tuba-seq^Ultra^)^44^. To generate analogous Tuba-seq^Ultra^ lentiviral libraries with pairs of sgRNAs, we synthesized dual-sgRNA oligo libraries and cloned them through a circular intermediate prior to ligation into a barcoded lentiviral-U6^BC^/Cre backbone (**Figure 1a** and **Supplementary Figure 1a**). PCR amplification and high-throughput sequencing of the BC-sgRNA1-sgRNA2 region from bulk tumor-bearing lungs from mice with tumors initiated with these vectors captures information on tumor genotype and size.

**Figure 1.**
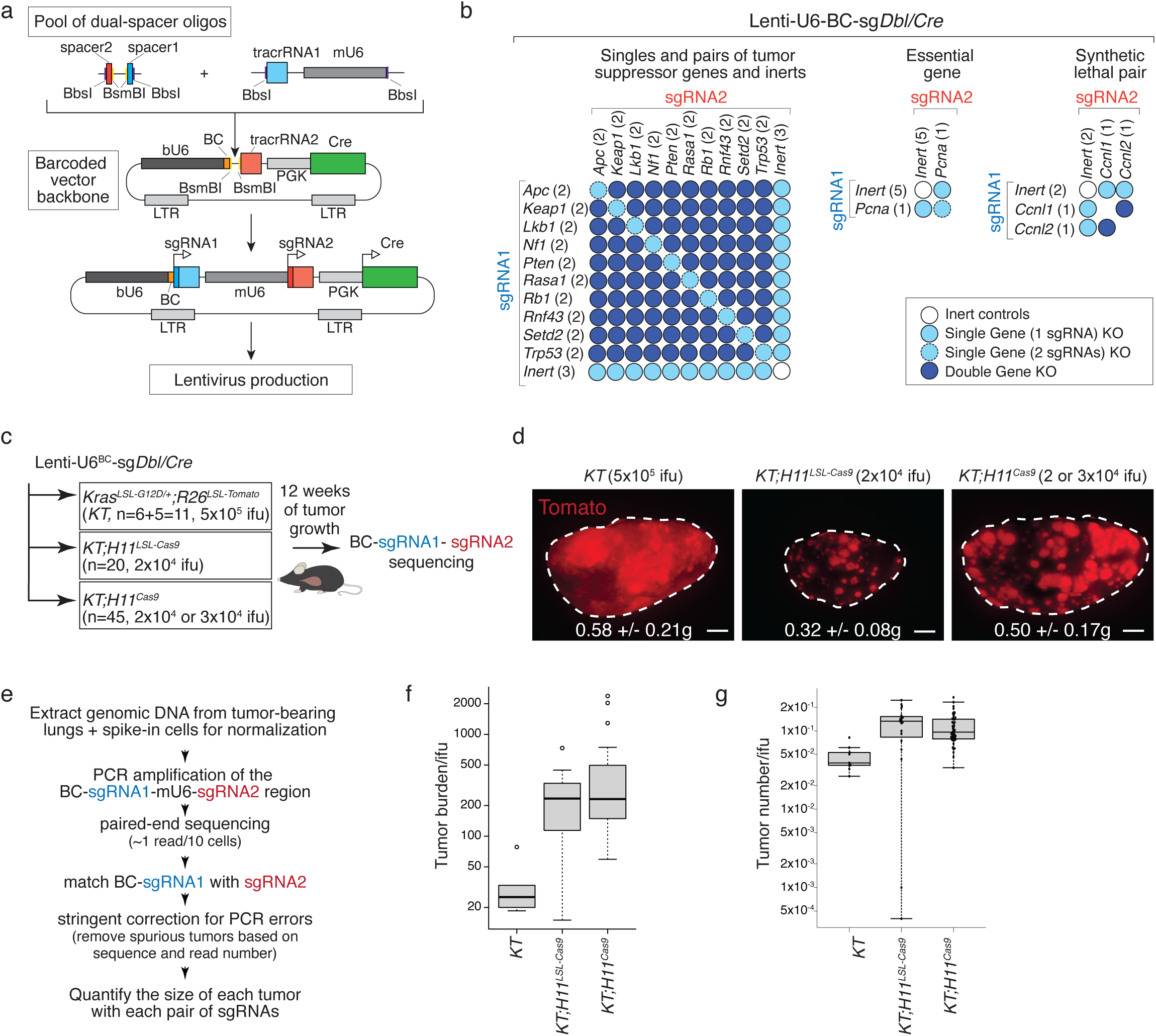
A dual-sgRNA system integrated with barcoding of clonal cell lineages and application to tumor suppressor gene epistasis in lung cancer. **a.** Generation of a library of dual-sgRNA vectors with an integrated diverse barcode to enable clonal tracking. **b.** sgRNA pairs in the Lenti-U6^BC^-sg*Dbl/Cre* pool. The sgRNA in position 1 (sgRNA1) and position 2 (sgRNA2) are indicated and the number of sgRNAs targeting each gene is shown in parentheses. This pool should create tumors with 45 distinct double knockout combinations. **c.** Initiation of KRAS^G12D^-driven lung tumors with Lenti-U6^BC^-sg*Dbl/Cre*. Genotype, lentiviral titer, and mouse number are indicated. Note that *Kras^LSL-G12D^;R26^LSL-Tomato^ (KT)* mice received 25-fold more virus than *KT;H11^LSL-Cas9^* and *KT;H11^LSL-Cas9^* mice. Bulk tumor-bearing lungs were analyzed after 12 weeks of tumor growth. **d.** Tomato fluorescent images of lung lobes from the indicated genotypes of mice. Mean +/- SD lung weight indicated. Scale bar, 2 mm. **e.** Outline of BC-sgRNA1-sgRNA2 read process, filtering, and analysis **f.** Box plot of tumor burden (total number of neoplastic cells of any genotype) normalized to lentiviral titer (ifu) across mouse genotype. **g.** Box plot of total number of tumors (any genotype of tumor estimated to have >50 neoplastic cells) normalized to lentiviral titer (ifu) across mouse genotype.

### Efficient inactivation of genes and gene pairs in somatic cells

To make use of this approach to study epistasis during tumorigenesis, we selected ten tumor suppressor genes that are commonly altered in lung adenocarcinoma and generated a dual-sgRNA oligo library targeting every pairwise combination of these genes with two sgRNAs each (Lenti-U6^BC^-sg*Dbl/Cre*; **Figure 1b, Supplementary Figure 1b**, and **Supplementary Table 1**). For comparison with the effects of single gene knockout, three Inert sgRNAs were paired with each tumor suppressor gene-targeting sgRNA. To assess Cas9 cutting efficiency from the first and second sgRNA positions within the vector, every sgRNA pair was created with both arrangements. Finally, the Lenti-U6^BC^-sg*Dbl/Cre* pool also contained an sgRNA targeting the essential gene *Pcna* paired with 5 Inert sgRNAs, as well as an sgRNA pair targeting the synthetic lethal genes *Ccln1* and *Ccln2* (**Figure 1b**)^24^.

We delivered the Lenti-U6^BC^-sg*Dbl/Cre* pool through intratracheal intubation to the lungs of *Kras^LSL-G12D/+^;Rosa26^LSL-Tomato^* (*KT*; n=11), *KT;H11^LSL-Cas9^* (n=20) and *KT;H11^Cas9^* (n=45) mice (**Figure 1c**). In these mice, the integrated lentiviral vector uniquely tags each transduced cell and the resulting tumors with a unique BC-sgRNA1-sgRNA2 element, Cre activates expression of oncogenic KRAS to initiate neoplastic growth, and Cre-mediated or constitutive Cas9 targets the genes defined by the sgRNAs. 12 weeks after lentiviral transduction, *KT;H11^LSL-Cas9^*and *KT;H11^Cas9^* mice had similar lung weights to *KT* mice despite receiving 25-fold less virus and had visibly larger tumors, consistent with dramatically increased growth of tumors that have combinations of tumor suppressor genes inactivated (**Figure 1d**).

The BC-sgRNA1-sgRNA2 region was PCR-amplified from genomic DNA extracted from tumor-bearing lungs followed by high-throughput sequencing (**Figure 1e** and **Supplementary Figure 1c**). Stringent filtering allowed us to quantify the number of reads from each clonal tumor with each pair of sgRNAs, while accurately excluding spurious reads generated by PCR uncoupling and sequencing errors (**Figure 1e, Supplementary Figure 1c**, and **Methods**)^45^. We used the number of reads from each tumor to calculate the number of neoplastic cells in each tumor, the number of tumors of each genotype, and several metrics of tumor growth (**Supplementary Figure 1d**). Analysis of tumors from *KT* mice showed that all sgRNA1-sgRNA2 pairs were well-represented in the pool (**Supplementary Figure 1e**).

The total tumor burden (defined as the total number of neoplastic cells of any genotype) normalized to lentiviral titer was about 10-fold higher in *KT;H11^LSL-Cas9^* and *KT;H11^Cas9^* mice relative to *KT* mice, consistent with increased tumor growth enabled by some genotypes (**Figure 1f**). *KT;H11^LSL-Cas9^* and *KT;H11^Cas9^* mice also had more tumors than *KT* mice when normalized to lentiviral titer, suggesting that inactivation of certain genes or gene pairs promotes tumor initiation and/or very early tumor growth and survival (**Figure 1g**).

Homozygous inactivation of the essential gene *Pcna* and homozygous inactivation of both *Ccln1* and *Ccln2* should reduce tumor number. We initially evaluated the determinants of efficient Cas9-mediated gene inactivation by quantifying the number of tumors with sg*Pcna* relative to sg*Inert*. Targeting *Pcna* reduced the number of tumors more in *KT;H11^Cas9^* mice than in *KT;H11^LSL-Cas9^* mice (**Figure 2a**). sg*Pcna* encoded in position 1 reduced tumor number modestly but consistently more than when encoded in position 2 in *KT;H11^LSL-Cas9^* mice, but both positions were similarly effective in *KT;H11^Cas9^* mice (**Figure 2a**). sg*Pcna* reduced tumor number similarly regardless of whether the sg*Inert* partner was Safe-targeting, non-targeting (NT and eGFP), or not expressed (NE), suggesting that sgRNA competition for Cas9 is not a major factor in driving target gene inactivation in this system (**Figure 2b**)^46^. Coincident targeting of *Ccnl1* and *Ccnl2*, but not targeting either gene individually, reduced tumor number specifically in the more efficient *KT;H11^Cas9^* background (**Figure 2c**). These data from our control vectors document highly efficient gene inactivation.

**Figure 2.**
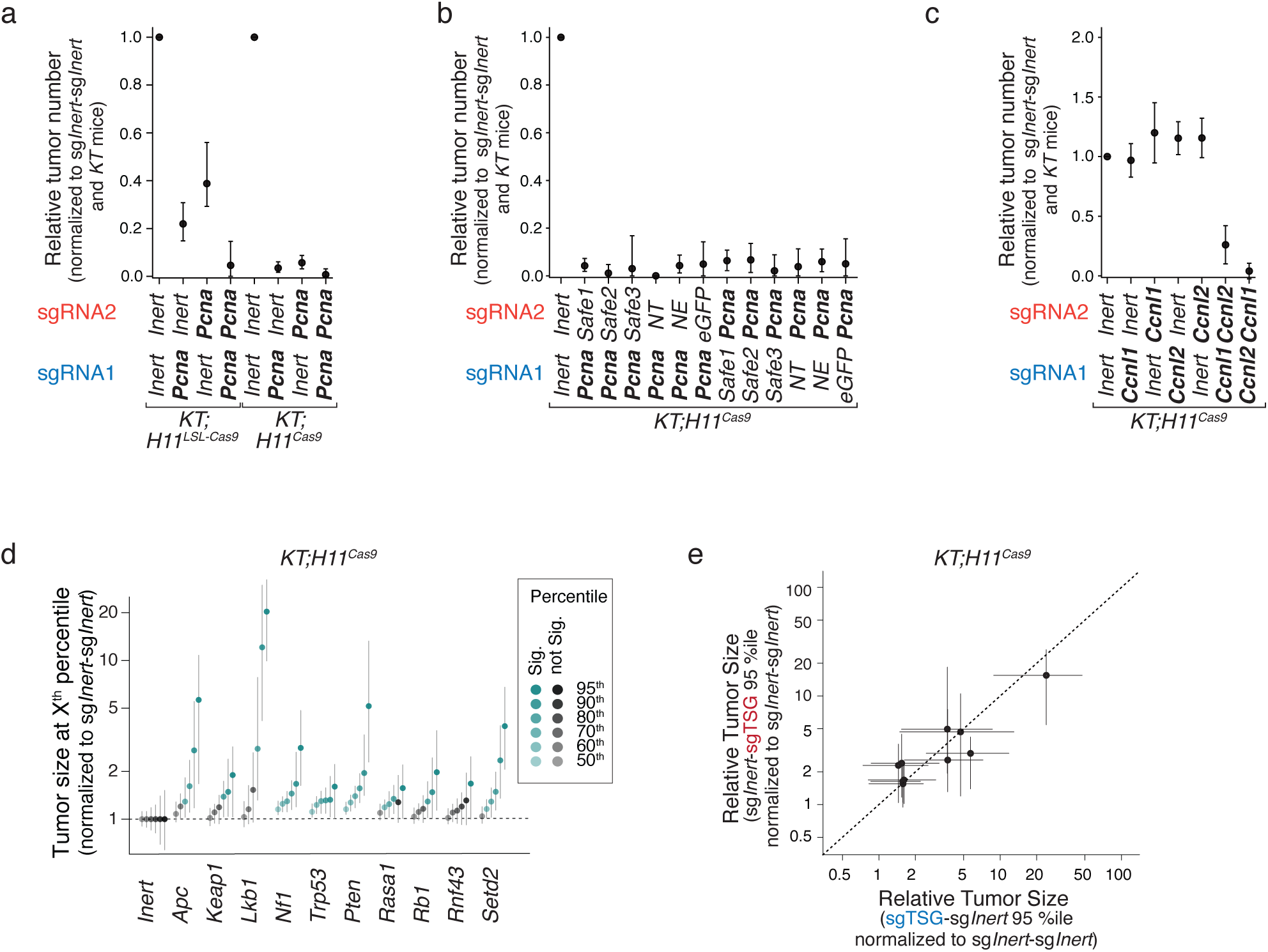
Efficient inactivation of an essential gene, a synthetic lethal pair, and tumor suppressor genes. **a.** Impact of targeting of an essential gene (*Pcna*) on tumor number in *KT;H11^LSL-Cas9^* and *KT;H11^Cas9^* mice with sg*Pcna* as sgRNA1 or sgRNA2 in the vector. Tumor number is normalized to the representation of each vector in *KT* mice (which lack Cas9) as well as to all sg*Inert*-sg*Inert* vectors. Mean +/- 95% confidence intervals are shown. **b.** Impact of targeting *Pcna* on tumor number in *KT;H11^Cas9^*mice across all vector configurations with sg*Pcna* as sgRNA1 or sgRNA2 in the vector and different non-targeting (*NT*), not expressed (*NE*), and *Safe*-targeting sg*Inerts*. sg*eGFP* is a non-targeting sg*Inert* that was previously suggested to be highly competative for Cas9. Mean +/- 95% confidence intervals are shown. **c.** Impact of targeting *Ccln1* and/or *Ccln2* (synthetic lethal pair) on tumor number in *KT;H11^Cas9^* mice across all vector configurations. Mean +/- 95% confidence intervals are shown. **d.** Tumor sizes for tumors with inactivation of each single tumor suppressor gene (TSG). Data are an aggregate of all sgTSG-sgInert, sgInert-sgTSG and sgTSG-sgTSG vectors targeting each TSG. 95% confidence intervals are shown. **e.** Comparison of effects on tumor size with TSG-targeting sgRNAs as sgRNA1 or sgRNA2 in a vector. Each dot represents a gene. Mean +/- 95% confidence intervals are shown.

We also evaluated the efficiency of Cas9-mediated gene inactivation within our dual-sgRNA system by examining the effects of targeting tumor suppressor genes with one sgRNA in either position or with two sgRNAs. Inactivation of individual tumor suppressor genes largely recapitulated previous data, with excellent resolution to identify effects on tumor size (**Figure 2d** and **Supplementary Figure 2a**) and results consistent between both sgRNA versions used to target each gene (**Supplementary Figure 2b-c**). In most cases, tumor suppressor gene-targeting sgRNAs did not have markedly different effects when encoded in position 1 compared to position 2 (**Figure 2e** and **Supplementary Figure 2d-e**). Targeting the same tumor suppressor gene with two sgRNAs instead of one modestly increased the effect in some cases, but for most tumor suppressor genes had no additional effect on tumor growth (**Supplementary Figure 2f**). Collectively, these data indicate that this barcoded dual-sgRNA system can efficiently inactivate genes in somatic cells *in vivo*, enabling a quantitative dissection of genetic interactions that influence tumor fitness.

### Inactivation of most tumor suppressor gene pairs shows diminishing returns epistasis

To identify epistatic interactions between pairs of tumor suppressor genes, we calculated the fitness of each tumor genotype (single and double mutant) relative to tumors with inert sgRNAs (**Figure 3a**). We compared the expected fitness of double mutants (the product of the relative fitness of both single mutants) to the actual observed double mutant fitness to calculate an epistasis score for each gene pair. Positive epistasis scores indicate synergistic effects and negative epistasis scores indicate buffering effects on tumor fitness (**Figure 3a-b**). To maximize the number of tumors supporting the interactions between each pair of tumor suppressor genes, we combined the data from the tumors with all sgRNAs targeting each gene as well as with the sgRNAs in either position. While the actual and expected fitness values for double mutants were highly correlated (**Supplementary Figure 3a**), most epistasis scores were negative (**Figure 3b-c**), suggesting that combined tumor suppressor gene inactivation generally has diminishing returns on tumor fitness. As expected, no epistatic effects were observed in *KT* mice lacking Cas9, in which no genes were inactivated (**Supplementary Figure 3b**). Moreover, epistasis scores showed a negative correlation with the expected fitness of a double mutant (**Figure 3c**), consistent with greater diminishing returns when single mutant genotypes already had high tumor fitness. When genes were clustered according to their epistasis scores across all gene pairs, *Apc* clustered separately from the rest of the genes, which otherwise formed two distinct clades (**Figure 3d**). Thus, while double tumor suppressor gene mutants still had higher fitness than their single mutant constituents, double mutants mostly had less-than-multiplicative effects, with a greater drop-off for high fitness single mutants.

**Figure 3.**
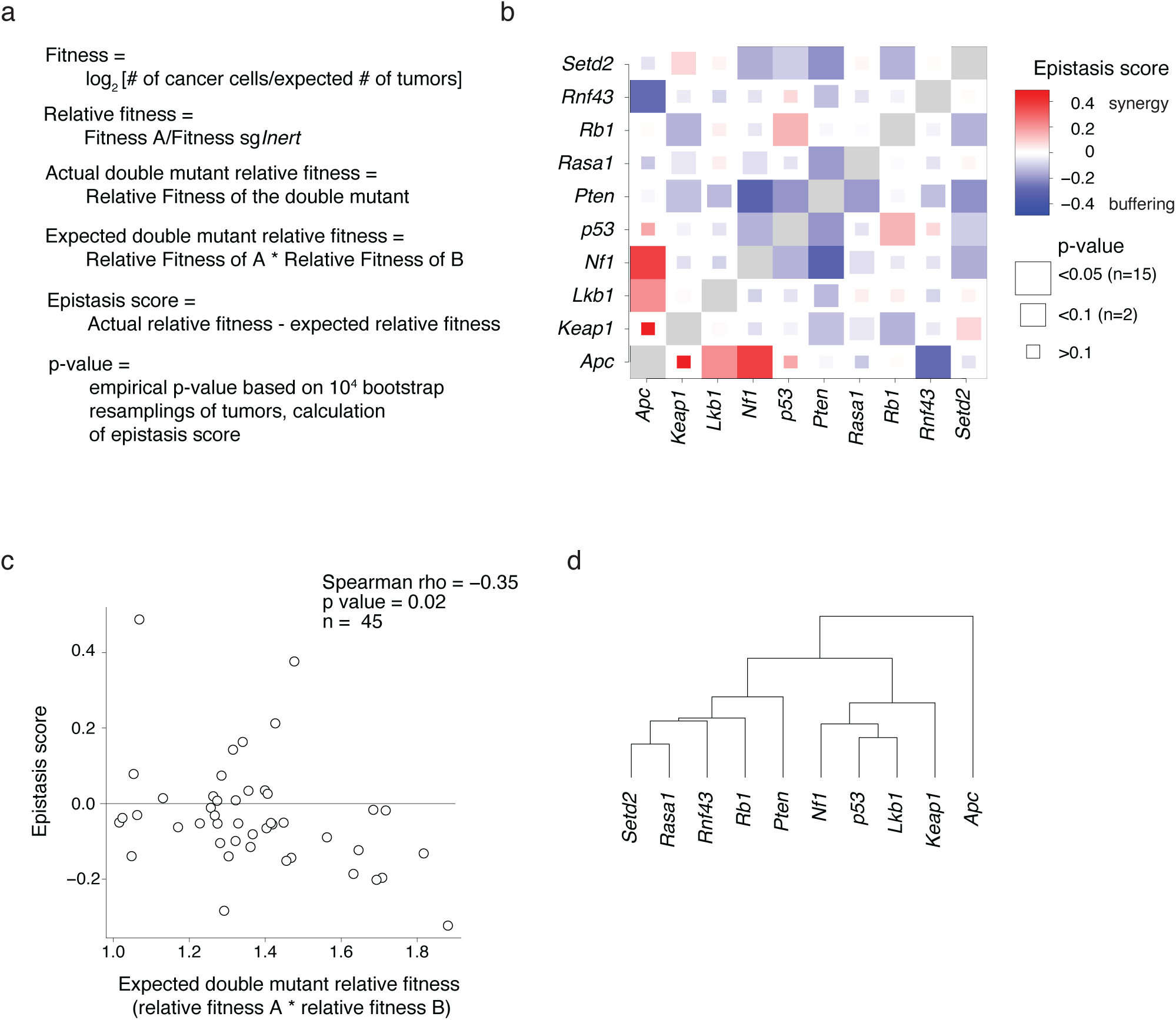
Broad Epistasis between tumor suppressor genes impacts lung cancer growth. **a.** Definitions and calculations of fitness and epistasis metrics. **b.** Heatmap of epistasis scores for all pairwise comparisons in *KT;H11^Cas9^* mice. The color indicates postive or negative epistasis and the box size indicates significance. The number of pairs with significant (p<0.05) and trending (p<0.1) positive and negative epistasis scores is indicated. **c.** Expected double mutant relative fitness and epistasis score negatively correlate. Each dot is a tumor suppressor gene pair. Spearman’s rho and associated p-value are indicated. **d.** Clustering of genes based on their epistasis scores across all shared pairs.

### *Apc* gene pairs are enriched for synergistic effects while *Pten* gene pairs most often have buffering effects

To explore gene-specific trends in epistasis, we calculated average epistasis values for every pair involving each gene (**Supplementary Figure 3c**). While most genes had slightly negative epistasis scores on average (consistent with overall trends), *Apc* stood out for having a positive average epistasis score (**Supplementary Figure 3c**). This was driven by synergistic interactions between inactivation of *Apc* and *Keap1*, *Lkb1*, *Nf1* and *p53* (**Figure 4a**), which had the highest epistasis scores across all data (**Supplementary Figure 3d**). In contrast, *Pten* exhibited strongly negative epistasis overall (**Figure 4b**), in keeping with the strong effect of *Pten* loss alone on tumor fitness (**Supplementary Figure 3c**).

**Figure 4.**
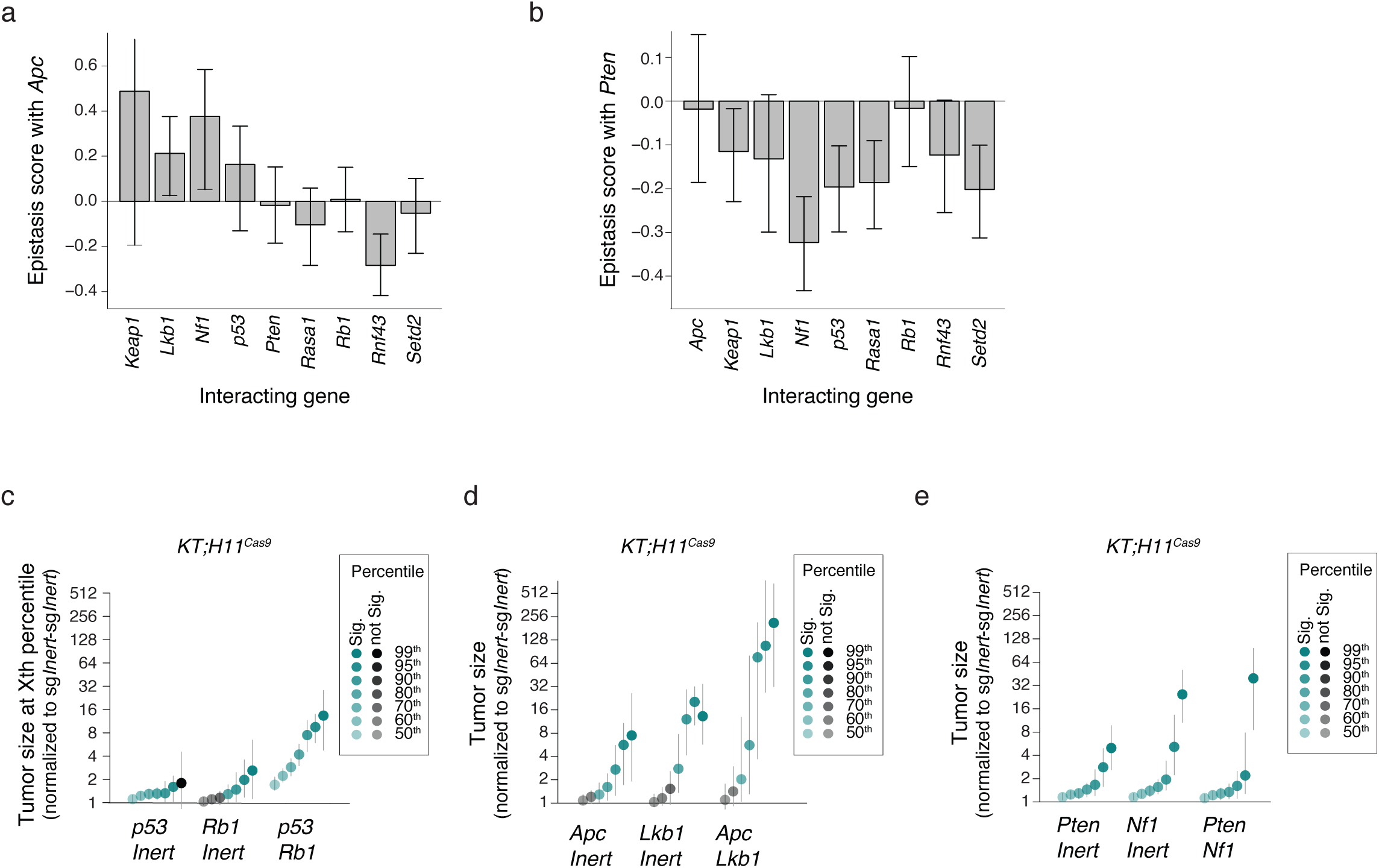
Tumor suppressor gene-specific genetic interactions influence tumor fitness. **a-b.** Epistasis scores +/- 95 percent confidence intervals for all pairs with Apc (**a**) and Pten (**b**). **c-e.** The size of tumors with each single and double mutant for *p53/Rb1* (**c**), *Apc/Lkb1* (**d**), and *Pten/Nf1* (**e**) at the given percentiles of the tumor size distribution normalized to sg*Inert*-sg*Inert* tumors. 95% confidence intervals are shown.

To uncover novel genetic interactions, we rank-ordered epistasis scores and assessed their statistical significance. A positive genetic interaction between *p53* and *Rb1* is well-established across cancer types, including being supported by mutational co-occurrence rates in human lung adenocarcinoma and by data from analogous mouse models of oncogenic Kras-driven lung adenocarcinoma^22,47–51^. Consistent with this, inactivation of *p53* and *Rb* had positive epistasis and a synergistic effect on tumor growth (**Figure 4c** and **Supplementary Figure 3d**). This confirms our ability to detect these types of synergistic genetic interactions and underscores the relative strength of the relationship between the P53 and RB pathways. *Apc-Lkb1* and *Apc-Nf1* had the most significant positive epistasis scores, with marked synergistic effects on tumor growth. Inactivation of these pairs resulted in the growth of exceptionally large tumors not created by any of the individual gene mutants (**Figure 4d**, **Supplementary Figure 4a**). Conversely, combined loss of *Pten* and *Nf1* had the most negative epistasis score among all gene pairs (**Supplementary Figure 3d**), with *Pten-Nf1* double mutant tumors being of a similar size to *Pten* or *Nf1* single mutant tumors (**Figure 4e**). As anticipated, no changes in tumor size were observed in *KT* mice lacking Cas9 (**Supplementary Figure 4b-e**). Thus, our quantitative approach for examining the effects of tumor genotype on fitness uncovered both strongly synergistic and buffering genetic interactions.

## DISCUSSION

Given the complexity of human tumor genotypes, understanding genetic interactions is critical to making sense of how tumor genotype drives critical aspects of tumorigenesis and tumor progression. In this study, we combined the pairwise inactivation of ten commonly mutated tumor suppressor genes with quantitative tumor barcoding and sequencing methods to deconvolute how these genes cooperate to promote tumor growth. While we observed several surprising synergistic interactions involving *Apc*, we found that the overwhelming majority of gene pairs showed diminishing returns, which could be the result of several factors. There is likely some degree of overlap in the pathways through which these tumor suppressor genes constrain tumor growth. *Nf1* and *Pten*, for instance, which had the strongest negative epistatic interaction in our data, are involved in RAS and PI3K signaling, respectively, two very closely linked pathways regulating cell proliferation and survival^52–55^. Similarly, *Pten* and the RAS suppressor *Rasa1* show negative epistasis, as do *Rasa1* and *Nf1*. Moreover, *Apc* and *Rnf43* both regulate the Wnt pathway and show strongly negative epistasis in our results^56–58^. Redundancy between or within pathways could therefore result in diminishing returns on tumor fitness. Furthermore, there is an upper limit to the degree to which cell proliferation can be increased through gene inactivation. As these tumors express oncogenic *Kras^G12D^*, they already have RAS pathway activation, which drives proliferation. As a result, they may have relatively little room to increase their proliferation even further compared to tumors without such a strong oncogenic driver. Concordantly, we previously showed that the combined inactivation of *Nf1*, *Rasa1* and *Pten* was highly synergistic in oncogene-negative lung adenocarcinoma^39,40^.

It is worth noting that the negative epistasis we observe experimentally is not synthetic lethality, in which an additional mutation reduces absolute tumor fitness compared to a single mutant. The double mutants that we investigated still increased overall tumor fitness, so mutations in these genes might still be expected to co-occur in human lung adenocarcinomas. Still, this implies that the exact functional impact of genetic interactions on tumor fitness cannot be inferred solely from tumor mutational data. Indeed, there are a number of confounding factors in uncovering genetic interactions from human data alone, including tumor subtype-specific mutations, variable tumor mutational burden among subtypes, as well as differing therapies and therapy resistance responses^19^.

The potential future applications of this platform in investigating tumorigenesis are multifold. It can be immediately applied to understand the effect of larger, more complete, pairwise combinations of tumor suppressors in lung adenocarcinoma. Lentiviral vectors with pairs of sgRNAs could also be combined with floxed alleles to create tumors with combinatorial inactivation of three genes, which may even better approximate the genetic makeup of human lung tumors. These quantitative and combinatorial approaches could be combined with CRISPRa, CRIPSRi, or Cas9-based methods for epigenetic modulation to increase or decrease the expression of pairs of genes *in vivo*. Moreover, this platform is not limited to lung adenocarcinoma and could be adapted to study other cancer types that can be initiated with viral vectors, including pancreatic cancer, bladder cancer, sarcoma, breast cancer, other types of lung cancer, and prostate cancer. This method could also contribute to the investigation of other aspects of tumor growth, progression, and metastasis. We defined epistasis effects based on tumor fitness, a metric derived largely from tumor size. However, given our data-rich readout, it should also be possible to describe genotypes based on other phenotypes, such as tumor initiation rate and the frequency of rare large tumors^35^. Creating more sophisticated models for tumor growth and epistasis across these measures might offer new insights. Finally, combinatorial inactivation of tumor suppressor genes and drug targets could uncover genotype-specific therapeutic effects, which could ultimately lead to more personalized patient treatments^37^.

These experiments have established and validated a platform for the rapid and quantitative analysis of the effect of combinatorial genetic alterations on tumor growth *in vivo*. These approaches have the potential to accelerate our understanding of the combinatorial driving forces of tumorigenesis.

## METHODS

### Generation of barcoded dual-sgRNA lentiviral vector libraries

To generate barcoded vector backbones, a 98 bp oligo containing a 16-nucleotide degenerate barcode (BC) and two BsmBI Type IIS restriction enzyme sites was cloned into the 3’ end of the bovine U6 promoter using Gibson Assembly (NEBuilder HiFi, NEB) (see **Supplementary Figure 1a**), in a vector also containing a tracrRNA (for guide 2) and PGK-Cre recombinase^44^. To create pooled dual-sgRNA inserts, designed combinations of guides were ordered as a single-stranded DNA pool (Twist Biosciences) of 138 bp oligos, each containing two guide spacers flanked by BsmBI and BbsI Type IIS restriction enzyme sites (all oligo sequences in **Supplementary Table 1**). Following low-cycle PCR amplification, these double guide oligos were ligated via Golden Gate Assembly (using BbsI) with a donor sequence containing a tracrRNA (for guide 1) and the mouse U6 promoter, resulting in a 528 bp circular intermediate. After PCR amplification to create linearized insert sequences, the inserts were cloned via Golden Gate Assembly (using BsmBI) into the barcoded vector backbone to yield the final pool of barcoded dual-sgRNA vectors (Lenti-U6^BC^-sg*Dbl/Cre*). This pooled product was then electroporated into competent E. Coli (C3020K, New England Biosciences) and plated onto LB-ampicillin plates. To ensure sufficient barcode diversity for Tuba-seq^Ultra^ sequencing and delineation of tumors, ∼10^6^ colonies were collected for the Lenti-U6^BC^- sg*Dbl/Cre* pool. This cloning procedure thereby enables the generation of lentiviral vectors containing any desired combinations of two sgRNAs in a highly multiplexed manner.

### Lentiviral packaging

To generate lentivirus, Lenti-U6^BC^-sg*Dbl/Cre* vectors were transfected as a pool into 293T cells with pCMV-VSV-G (Addgene #8454) envelope plasmid and pCMV-dR8.2 dvpr (Addgene #8455) packaging plasmid using polyethylenimine (Polysciences). Sodium butyrate (Sigma Alrich, B5887) was added 8 hours after transfection to a final concentration of 20 mM to inhibit silencing of viral expression, and medium was changed after 24 hours. Supernatants were collected 36 and 48 hours after transfection, filtered with 0.45 µm syringe filters (Millipore, SLHP033RB) to remove cells and cell debris, concentrated by ultracentrifugation (25,000 g for 1.5 hours at 4°C) and resuspended in PBS overnight. Each virus was titered against a known standard using LSL-YFP Mouse Embryonic Fibroblasts (MEFs) (a gift from Dr. Alejandro Sweet-Cordero/UCSF). Lentiviral vector aliquots were stored at -80°C and were thawed and pooled immediately prior to delivery to mice.

### Animal Studies

The use of mice for this study was approved by the Institutional Animal Care and Use Committee at Stanford University, protocol number 26696. *Kras^LSL-G12D/+^*(RRID:IMSR_JAX:008179)^59^, *R26^LSL-tdTomato^* (RRID:IMSR_JAX:007914)^60^, *H11^LSL-Cas9^* (RRID:IMSR_JAX:027632)^61^ and *H11^Cas9^* mice (RRID:IMSR_JAX:006054) mice have been previously described. All mice were on a pure C57BL/6 background.

Tumors were initiated by intratracheal delivery of 60 ul of resuspended lentiviral vectors in PBS. Lentiviral titer and time of tumor development are indicated in each figure.

### **T**umor Barcode Sequencing and Analysis

Tuba-seq^Ultra^ libraries were generated by isolating genomic DNA from bulk tumor-bearing lung tissue, followed by PCR amplification of the BC-sgRNA1-sgRNA2 region from 32 μg of bulk lung genomic DNA using Q5 Ultra II High-Fidelity 2× Master Mix (New England Biolabs, M0494X). Unique dual-indexed primers were used to amplify each sample followed by purification using Agencourt AMPure XP beads (Beckman Coulter, A63881). The libraries were pooled based on lung weights to ensure even read depth and sequenced (read length 2 × 150 bp) on the Illumina NextSeq 6000 platform. Tuba-seq analysis of tumor barcode reads was performed as previously described^34,35^, except that sgRNAs were first directly sequenced and their sequences matched with our intended target genes and guide versions (e.g. Apc-V1). At library preparation, a template switching event might happen between the two sgRNA regions. This can create chimera sequences, groups of BC-sgRNA1-sgRNA2 regions that are created from two distinct tumors via PCR template switching. The resulting sequencing reads of groups of BC-sgRNA1-sgRNA2 do not correspond to any real tumor. With the diverse barcode region and a matching sgRNA1 sequence close to the DNA-barcode, we can easily remove these PCR artifacts by simply removing the “tumors” with the lower read count from all such pairs.

### Calculation of Relative Fitness of Each Genotype of Tumors

Fitness of each tumor genotype was approximated by the growth rate of tumors. Fitness was calculated for each tumor genotype in a given mouse strain by considering the expected number of cells with each lentiviral vector at the start of the experiment (#start*_genotype_mouse-strain_*) and number of neoplastic cells with each lentiviral vector after tumor growth at the end of the experiment (#end*_genotype_mouse-strain_*).

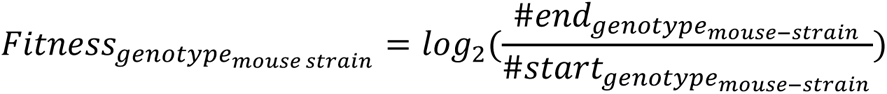

#end*_genotype_mouse-strain_* is the sum of all neoplastic cells in all tumors of that genotype at the end of the experiment from all mice of a given strain.

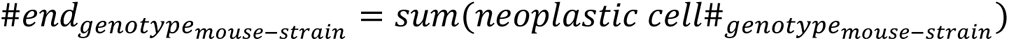

#start*_genotype_mouse-strain_* is the expected number of cells of that genotype at the start of the experiment. #start*_genotype_mouse-strain_* was determined from the number of tumors with each viral vector (titer*_genotype_*) from *KT* Cas9-negative control mice. In *KT* mice, the number of tumors with each vector represents the exact titer of each vector (titer_genotype_) in the lentiviral pool. Data from *KT* mice were used to calculate the relative titer across genotypes. To put #start_genotype_ in the context of the most potent genotype (resulting in the most tumors per titer units) in the given mouse strain, we normalized the number of tumors with each lentiviral vector in *KT* mice (titer_genotype_) to the number of *Nf1;Pten* double mutant tumors in *KT* mice (titer*_Nf1;Pten_*) and then multiplied this by the total number of *Nf1;Pten* double mutant tumors across all mice of a given strain (TumorNumber*_Nf1;Pten_mouse-strain_*). Thus, #start*_genotype_mouse-strain_* represents the titer-corrected expected number of tumors of that genotype if that genotype were as potent as double *Nf1;Pten* mutation in that strain. As an example, #start for single mutant genotype *Apc* in *KTC* mice (#start *_Apc_KTC_*) was calculated as:

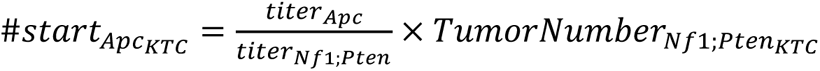

Finally, we calculated “Relative Fitness” within each mouse strain by normalizing the fitness of each genotype to the fitness of tumors with the inert sgRNAs in the lentiviral vector.

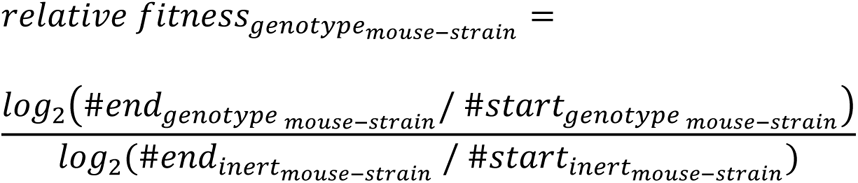

The above defined measure is a commonly used fitness definition^37,62–65^.

### Statistical Analysis

All statistical analyses were performed using the R software environment. P values and 95% CIs (represented by whiskers) were calculated using bootstrap resampling (10,000 repetitions). Bootstrapping was done by random resampling with replacement of all the tumors in all of the mice of a given strain. For **Figure 2a-e**, **Figure 4c-e**, **Supplementary Figure 2a-f**, and **Supplementary Figure 4a-e**, a nested bootstrap approach was used where first all the mice from a given strain were resampled with replacement, and then, all tumors in this virtual mouse cohort were resampled again with replacement.

## ACKNOWLEDGEMENTS

We thank the Stanford Veterinary Animal Care Staff for expert animal care and members of the Winslow laboratory for helpful comments. J.D.H was supported by an American Cancer Society postdoctoral fellowship (PF-21-112-01-MM) and a Tobacco-Related Disease Research Program (TRDRP) fellowship (T31FT1619). G.B. was supported by a TRDRP Fellowship (T31FT-1619), by the National Research, Development and Innovation Office in Hungary (RRF-2.3.1-21-2022-00006), and by the HUN-REN Welcome Home and Foreign Researcher Recruitment Programme (KSZF-139/2023). This work was supported by NIH R01-CA230025 (to M.M.W), NIH R01-CA231253 (to M.M.W and D.A.P), NIH R01-CA234349 (to M.M.W and D.A.P.), and in part by the Stanford Cancer Institute support grant (NIH P30-CA124435).

---

**Supplementary Figure 1.**
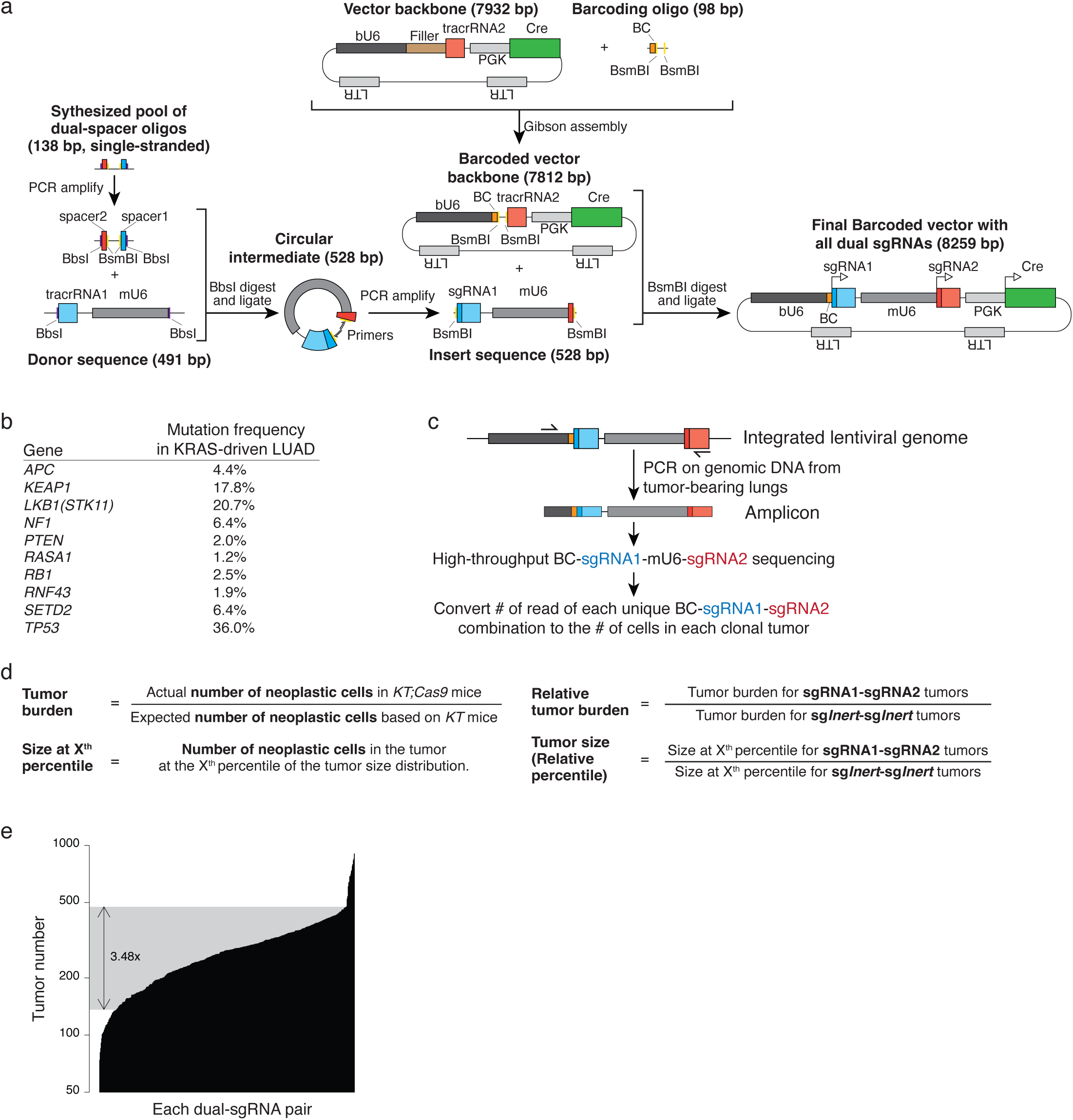
Generation of dual-sgRNA libraries with diverse barcodes to uniquely tag each clonal tumor lineage. **a.** Detailed outline of the cloning to generate the barcoded dual-sgRNA vector pool. **b.** Frequency of mutation of each tumor suppressor gene in this study. Data from lung adenocarcincinomas with KRAS mutations in GENIE (cohort v15.0). **c.** Overview of the Tuba-seq^Ultra^ pipeline to quantify tumor number (# of barcodes) and size (# of reads of each BC-sgRNA1-sgRNA2). **d.** Definitions of tumor burden and relative tumor size metrics which are used throughout this manuscript (see Methods). **e.** Representation of all dual-sgRNA pairs across all tumors within *KT* mice. Shaded area shows middle 90% of data (5th to 95th quantile), which represents a 3.48-fold difference in tumor number.

**Supplementary Figure 2.**
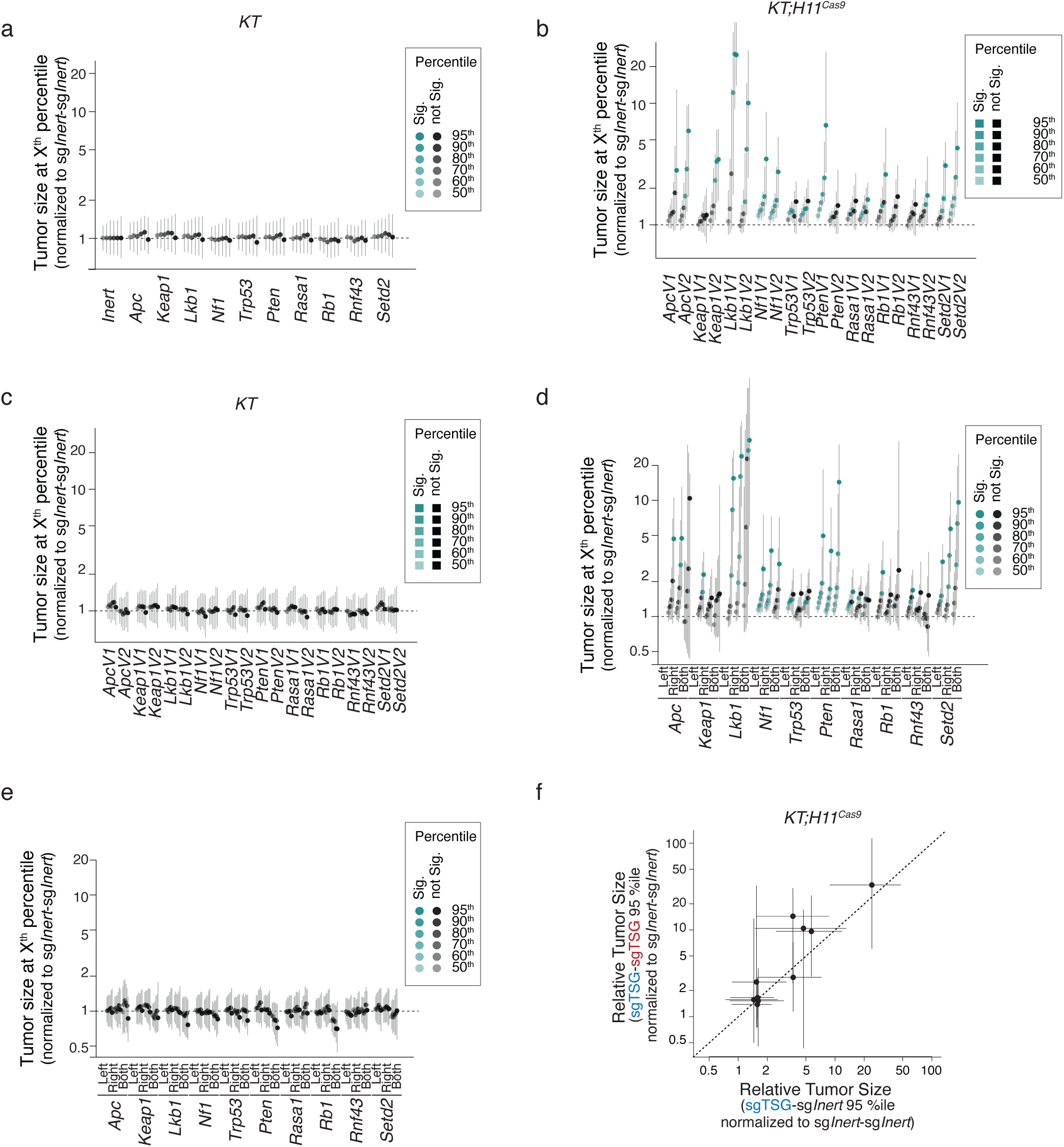
Tumor barcoding increases the resolution to find tumor suppressor effects. **a.** Tumor sizes for tumors with sgRNAs targeting each single tumor suppressor gene (TSG) in *KT* mice (without Cas9). Data are an aggregate of all sgTSG-sgInert, sgInert-sgTSG and sgTSG-sgTSG vectors targeting each TSG. 95% confidence intervals are shown. **b-c.** Tumor sizes in *KT;H11^Cas9^* (**b**) and *KT* (**c**) mice for different guide versions targeting each single TSG. 95% confidence intervals are shown. **d-e.** Tumor sizes in *KT;H11^Cas9^* (**d**) and *KT* (**e**) mice for tumors with single sgRNAs targeting each TSG from position 1 (left) or position 2 (right), as well as tumors targeting each TSG with two sgRNAs (both). 95% confidence intervals are shown. **f.** Comparison of effects on tumor size when targeting each TSG with one or two sgRNAs in a vector. Each dot represents a gene. Mean +/- 95% confidence intervals are shown.

**Supplementary Figure 3.**
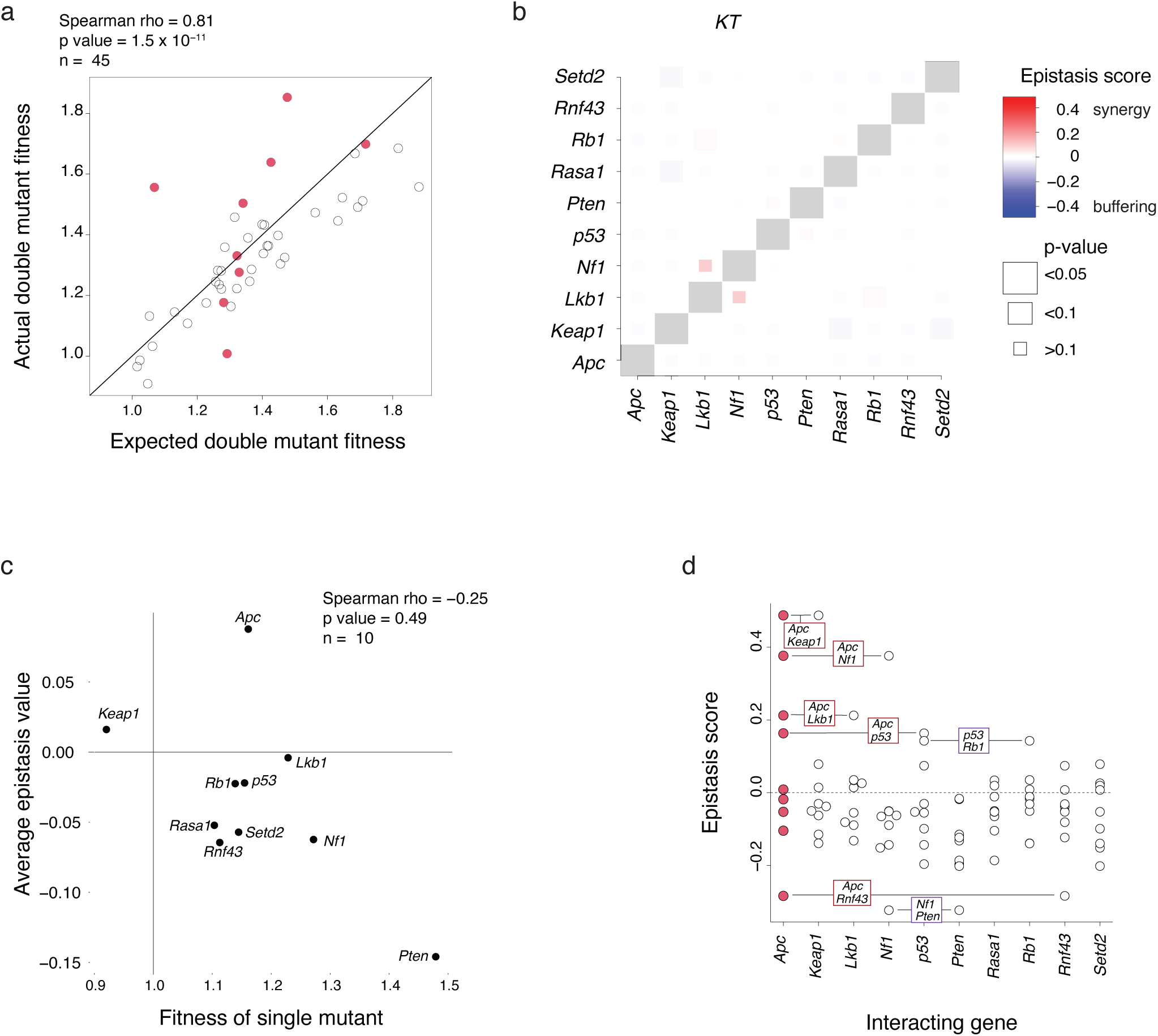
Epistasis between tumor suppressors drive lung cancer growth. **a.** Expected versus actual double mutant fitness for each of the 45 pairs of tumor suppressor genotypes, with pairs including *Apc* highlighted in red. Epistasis score is the difference between the actual and expected relative fitness values. Spearman’s rho and associated p-value are indicated. **b.** Heatmap of epistasis scores for all pairwise comparisons in *KT* mice (lacking Cas9). The color indicates postive or negative epistasis and the box size indicates significance. No pairs have significant epistasis scores. **c.** Average epistasis scores for all pairwise combinations involving the indicated gene versus the fitness of tumors with inactivation of that gene alone. Spearman’s rho and associated p-value are indicated. **d.** Individual epistasis scores for all pairwise combinations involving each gene. Scores are plotted for both genes in a given pair, so duplicate scores are present (indicated for key genetic interaction pairs).

**Supplementary Figure 4.**
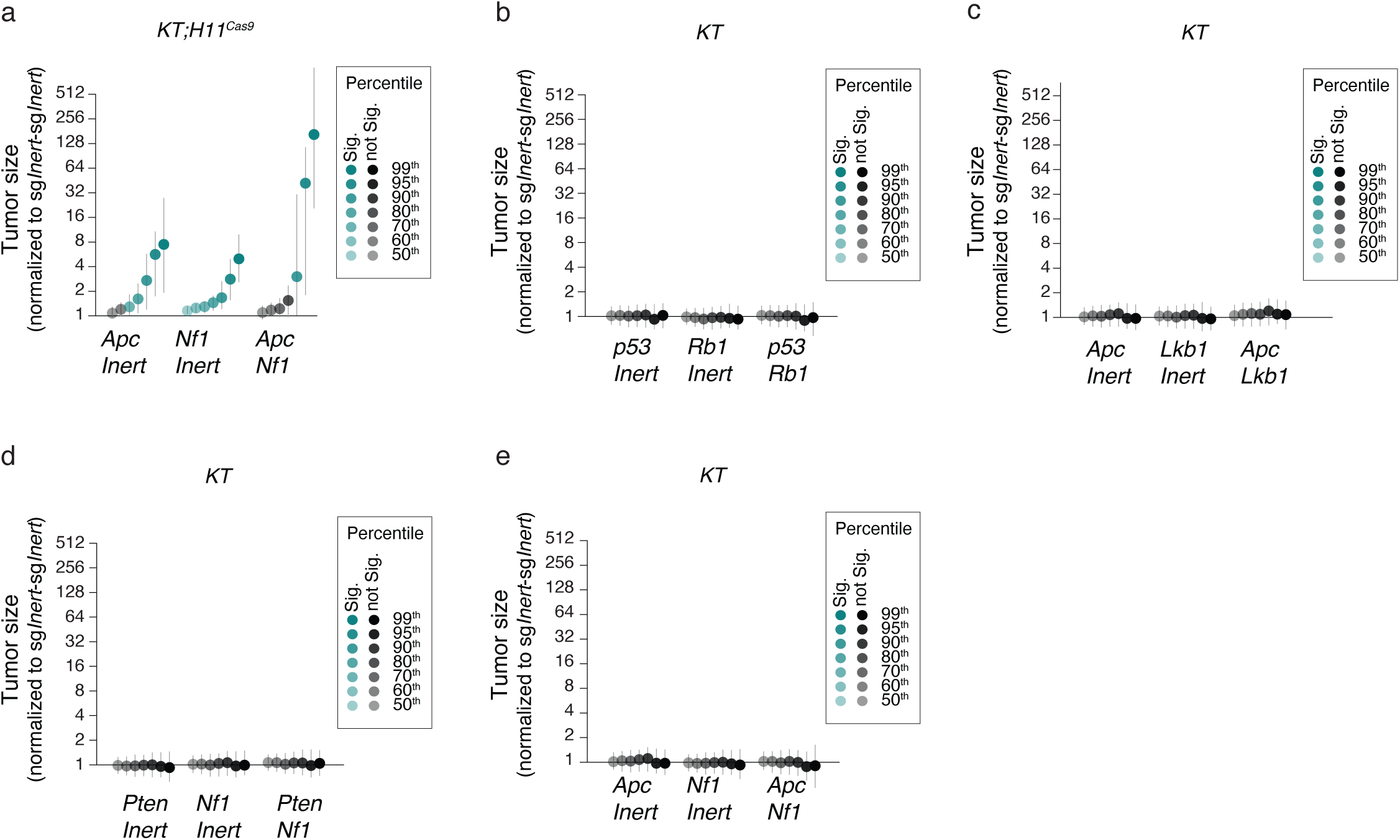
No effect of mice that lack Cas9 and tumor size data for a synergistic pair. **a-e.** The size of tumors with each single and double mutant for *Apc/Nf1* in *KT;H11^Cas9^* mice (**a**) *p53/Rb1* in *KT* mice (**b**), *Apc/Lkb1* in *KT* mice (**c**), *Pten/Nf1* in *KT* mice (**d**), and *Apc/Nf1* in *KT* mice (**e**) at the given percentiles of the tumor size distribution normalized to sg*Inert*-sg*Inert* tumors. 95% confidence intervals are shown.

## References

1. Boyle, E. A., Li, Y. I. & Pritchard, J. K. An Expanded View of Complex Traits: From Polygenic to Omnigenic. Cell 169, 1177–1186 (2017).

2. Civelek, M. & Lusis, A. J. Systems genetics approaches to understand complex traits. Nat Rev Genet 15, 34–48 (2014).

3. Seldin, M., Yang, X. & Lusis, A. J. Systems genetics applications in metabolism research. Nat Metab 1, 1038–1050 (2019).

4. Wray, N. R., Wijmenga, C., Sullivan, P. F., Yang, J. & Visscher, P. M. Common Disease Is More Complex Than Implied by the Core Gene Omnigenic Model. Cell 173, 1573–1580 (2018).

5. Jahchan, N. S. et al. A drug repositioning approach identifies tricyclic antidepressants as inhibitors of small cell lung cancer and other neuroendocrine tumors. Cancer Discov 3, 1364– 1377 (2013).

6. Jameson, K. L. et al. IQGAP1 scaffold-kinase interaction blockade selectively targets RAS-MAP kinase–driven tumors. Nature Medicine 19, 626–630 (2013).

7. Ehrenreich, I. M., Gerke, J. P. & Kruglyak, L. Genetic dissection of complex traits in yeast: insights from studies of gene expression and other phenotypes in the BYxRM cross. Cold Spring Harb Symp Quant Biol 74, 145–153 (2009).

8. VanderSluis, B. et al. Genetic interactions reveal the evolutionary trajectories of duplicate genes. Molecular Systems Biology 6, 429 (2010).

9. Costanzo, M. et al. The Genetic Landscape of a Cell. Science 327, 425–431 (2010).

10. Costanzo, M. et al. A global genetic interaction network maps a wiring diagram of cellular function. Science 353, aaf1420 (2016).

11. Winslow, M. M. et al. Suppression of lung adenocarcinoma progression by Nkx2-1. Nature 473, 101–104 (2011).

12. Mazur, P. K. et al. SMYD3 links lysine methylation of MAP3K2 to Ras-driven cancer. Nature 510, 283–287 (2014).

13. Xue, W. et al. Response and resistance to NF-κB inhibitors in mouse models of lung adenocarcinoma. Cancer Discov 1, 236–247 (2011).

14. Aaltonen, L. A. et al. Pan-cancer analysis of whole genomes. Nature 578, 82–93 (2020).

15. Nguyen, B. et al. Genomic characterization of metastatic patterns from prospective clinical sequencing of 25,000 patients. Cell 185, 563–575.e11 (2022).

16. Birkbak, N. J. & McGranahan, N. Cancer Genome Evolutionary Trajectories in Metastasis. Cancer Cell 37, 8–19 (2020).

17. Lee, M. C. et al. A multiplexed in vivo approach to identify driver genes in small cell lung cancer. Cell Rep 42, 111990 (2023).

18. Hebert, J. D., Neal, J. W. & Winslow, M. M. Dissecting metastasis using preclinical models and methods. Nat Rev Cancer 23, 391–407 (2023).

19. van de Haar, J. et al. Identifying Epistasis in Cancer Genomes: A Delicate Affair. Cell 177, 1375–1383 (2019).

20. Kandoth, C. et al. Mutational landscape and significance across 12 major cancer types. Nature 502, 333–339 (2013).

21. Sanchez-Vega, F. et al. Oncogenic Signaling Pathways in The Cancer Genome Atlas. Cell 173, 321–337.e10 (2018).

22. Rogers, Z. N. et al. Mapping the in vivo fitness landscape of lung adenocarcinoma tumor suppression in mice. Nat. Genet. 50, 483–486 (2018).

23. Foggetti, G. et al. Genetic determinants of EGFR-Driven Lung Cancer Growth and Therapeutic Response In Vivo. Cancer Discov (2021) doi:10.1158/2159-8290.CD-20-1385.

24. Parrish, P. C. R. et al. Discovery of synthetic lethal and tumor suppressor paralog pairs in the human genome. Cell Rep 36, 109597 (2021).

25. Zhao, X., Li, J., Liu, Z. & Powers, S. Combinatorial CRISPR/Cas9 Screening Reveals Epistatic Networks of Interacting Tumor Suppressor Genes and Therapeutic Targets in Human Breast Cancer. Cancer Res 81, 6090–6105 (2021).

26. Heitink, L. et al. In vivo genome-editing screen identifies tumor suppressor genes that cooperate with Trp53 loss during mammary tumorigenesis. Mol Oncol 16, 1119–1131 (2022).

27. Chiou, S.-H. et al. A conditional system to specifically link disruption of protein-coding function with reporter expression in mice. Cell Rep 7, 2078–2086 (2014).

28. Xue, W. et al. CRISPR-mediated direct mutation of cancer genes in the mouse liver. Nature 514, 380–384 (2014).

29. Platt, R. J. et al. CRISPR-Cas9 Knockin Mice for Genome Editing and Cancer Modeling. Cell 159, 440–455 (2014).

30. Yin, H. et al. Genome editing with Cas9 in adult mice corrects a disease mutation and phenotype. Nat Biotechnol 32, 551–553 (2014).

31. Walter, D. M. et al. RB constrains lineage fidelity and multiple stages of tumour progression and metastasis. Nature 569, 423–427 (2019).

32. Caswell, D. R. et al. Obligate Progression Precedes Lung Adenocarcinoma Dissemination. Cancer Discovery 4, 781–789 (2014).

33. Tang, Y. J., Shuldiner, E. G., Karmakar, S. & Winslow, M. M. High-Throughput Identification, Modeling, and Analysis of Cancer Driver Genes In Vivo. Cold Spring Harb Perspect Med 13, a041382 (2023).

34. Rogers, Z. N. et al. A quantitative and multiplexed approach to uncover the fitness landscape of tumor suppression in vivo. Nat. Methods 14, 737–742 (2017).

35. Cai, H. et al. A functional taxonomy of tumor suppression in oncogenic KRAS-driven lung cancer. Cancer Discov (2021) doi:10.1158/2159-8290.CD-20-1325.

36. Michlits, G. et al. CRISPR-UMI: single-cell lineage tracing of pooled CRISPR-Cas9 screens. Nat Methods 14, 1191–1197 (2017).

37. Han, K. et al. Synergistic drug combinations for cancer identified in a CRISPR screen for pairwise genetic interactions. Nature Biotechnology 35, 463–474 (2017).

38. Park, J. J. et al. Double knockout CRISPR screen for cancer resistance to T cell cytotoxicity. J Hematol Oncol 15, 172 (2022).

39. Yousefi, M. et al. Combinatorial Inactivation of Tumor Suppressors Efficiently Initiates Lung Adenocarcinoma with Therapeutic Vulnerabilities. Cancer Res 82, 1589–1602 (2022).

40. Yousefi, M. et al. Fully accessible fitness landscape of oncogene-negative lung adenocarcinoma. Proc Natl Acad Sci U S A 120, e2303224120 (2023).

41. Murray, C. W. et al. An LKB1-SIK Axis Suppresses Lung Tumor Growth and Controls Differentiation. Cancer Discov 9, 1590–1605 (2019).

42. Murray, C. W. et al. LKB1 drives stasis and C/EBP-mediated reprogramming to an alveolar type II fate in lung cancer. Nat Commun 13, 1090 (2022).

43. Hill, A. J. et al. On the design of CRISPR-based single-cell molecular screens. Nat Methods 15, 271–274 (2018).

44. Tang, Y. J. et al. A functional map of epigenetic regulators in lung carcinogenesis. [Manuscript in preparation].

45. Hegde, M., Strand, C., Hanna, R. E. & Doench, J. G. Uncoupling of sgRNAs from their associated barcodes during PCR amplification of combinatorial CRISPR screens. PLoS One 13, e0197547 (2018).

46. Najm, F. J. et al. Orthologous CRISPR-Cas9 enzymes for combinatorial genetic screens. Nat Biotechnol 36, 179–189 (2018).

47. Zhou, Z. et al. Synergy of p53 and Rb Deficiency in a Conditional Mouse Model for Metastatic Prostate Cancer. Cancer Research 66, 7889–7898 (2006).

48. Yamaguchi, T., Ikehara, S., Nakanishi, H. & Ikehara, Y. A genetically engineered mouse model developing rapid progressive pancreatic ductal adenocarcinoma. J Pathol 234, 228–238 (2014).

49. Dosaka-Akita, H. et al. Altered retinoblastoma protein expression in nonsmall cell lung cancer: its synergistic effects with altered ras and p53 protein status on prognosis. Cancer 79, 1329–1337 (1997).

50. Xu, H. J., Cagle, P. T., Hu, S. X., Li, J. & Benedict, W. F. Altered retinoblastoma and p53 protein status in non-small cell carcinoma of the lung: potential synergistic effects on prognosis. Clin Cancer Res 2, 1169–1176 (1996).

51. Xing, E. P., Yang, G. Y., Wang, L. D., Shi, S. T. & Yang, C. S. Loss of heterozygosity of the Rb gene correlates with pRb protein expression and associates with p53 alteration in human esophageal cancer. Clin Cancer Res 5, 1231–1240 (1999).

52. Nussinov, R., Tsai, C.-J. & Jang, H. Anticancer drug resistance: An update and perspective. Drug Resist Updat 59, 100796 (2021).

53. Cuesta, C., Arévalo-Alameda, C. & Castellano, E. The Importance of Being PI3K in the RAS Signaling Network. Genes (Basel) 12, 1094 (2021).

54. Krygowska, A. A. & Castellano, E. PI3K: A Crucial Piece in the RAS Signaling Puzzle. Cold Spring Harb Perspect Med 8, a031450 (2018).

55. Shaw, R. J. & Cantley, L. C. Ras, PI(3)K and mTOR signalling controls tumour cell growth. Nature 441, 424–430 (2006).

56. Bond, C. E. et al. RNF43 and ZNRF3 are commonly altered in serrated pathway colorectal tumorigenesis. Oncotarget 7, 70589–70600 (2016).

57. Yaeger, R. et al. Clinical Sequencing Defines the Genomic Landscape of Metastatic Colorectal Cancer. Cancer Cell 33, 125–136.e3 (2018).

58. van de Wetering, M. et al. Prospective derivation of a living organoid biobank of colorectal cancer patients. Cell 161, 933–945 (2015).

59. Jackson, E. L. et al. Analysis of lung tumor initiation and progression using conditional expression of oncogenic K-ras. Genes Dev. 15, 3243–3248 (2001).

60. Madisen, L. et al. A robust and high-throughput Cre reporting and characterization system for the whole mouse brain. Nat Neurosci 13, 133–140 (2010).

61. Chiou, S.-H. et al. Pancreatic cancer modeling using retrograde viral vector delivery and in vivo CRISPR/Cas9-mediated somatic genome editing. Genes Dev. 29, 1576–1585 (2015).

62. Remold, S. K. & Lenski, R. E. Contribution of individual random mutations to genotype-by-environment interactions in Escherichia coli. Proceedings of the National Academy of Sciences 98, 11388–11393 (2001).

63. Lenski, R. E., Rose, M. R., Simpson, S. C. & Tadler, S. C. Long-term experimental evolution in Escherichia coli. I. Adaptation and divergence during 2,000 generations. American Naturalist 138, 1315–1341 (1991).

64. Melnyk, A. H., Wong, A. & Kassen, R. The fitness costs of antibiotic resistance mutations. Evolutionary Applications 8, 273–283 (2015).

65. Alonso-del Valle, A. et al. Variability of plasmid fitness effects contributes to plasmid. Nat Commun 12, 2653 (2021).

